# A hierarchical Bayesian framework for understanding the spatiotemporal dynamics of the intestinal epithelium

**DOI:** 10.1101/072561

**Authors:** O.J. Maclaren, A. Parker, C. Pin, S.R. Carding, A.J.M. Watson, A.G. Fletcher, H.M. Byrne, P.K. Maini

## Abstract

Our work addresses two key challenges, one biological and one methodological. First, we aim to understand how proliferation and cellular migration rates in the intestinal epithelium are related under healthy, damaged (Ara-C treated) and recovering conditions, and how these relations can be used to identify mechanisms of repair and regeneration. We analyse new data, presented in more detail in a companion paper, in which BrdU/IdU cell-labelling experiments were performed under these respective conditions. Second, in considering how to more rigorously process these data and interpret them using mathematical models, we develop a probabilistic, hierarchical framework. This framework provides a best-practice approach for systematically modelling and understanding the uncertainties that can otherwise undermine drawing reliable conclusions - uncertainties in experimental measurement and treatment, difficult-to-compare mathematical models of underlying mechanisms, and unknown or unobserved parameters. Both discrete and continuous mechanistic models are considered and related via hierarchical conditional probability assumptions. This allows the incorporation of features of both continuum tissue models and discrete cellular models. We perform model checks on both in-sample and out-of-sample datasets and use these checks to illustrate how to test possible model improvements and assess the robustness of our conclusions. This allows us to consider - and ultimately decide against - the need to retain finite-cell-size effects to explain a small misfit appearing in one set of long-time, out-of-sample predictions. Our approach leads us to conclude, for the present set of experiments, that a primarily proliferation-driven model is adequate for predictions over most time-scales. We describe each stage of our framework in detail, and hope that the present work may also serve as a guide for other applications of the hierarchical approach to problems in computational and systems biology more generally.

**Author Summary:** The intestinal epithelium serves as an important model system for studying the dynamics and regulation of multicellular populations. It is characterised by rapid rates of self-renewal and repair; failure of the regulation of these processes is thought to explain, in part, why many tumours occur in the intestinal and similar epithelial tissues. These features have led to a large amount of work on estimating rate parameters in the intestine. There still remain, however, large gaps between the raw data collected, the experimental interpretation of these data, and speculative mechanistic models for underlying processes. In our view hierarchical statistical modelling provides an ideal, but currently underutilised, method to begin to bridge these gaps. This approach makes essential use of the distinction between ‘measurement’, ‘process’ and ‘parameter’ models, giving an explicit framework for combining experimental data and mechanistic modelling in the presence of multiple sources of uncertainty. As we illustrate, the hierarchical approach also provides a suitable framework for addressing other methodological questions of broader interest in systems biology: how to systematically relate discrete and continuous mechanistic models; how to formally interpret and visualise statistical evidence; and how to express causal assumptions in terms of conditional independence.

## Introduction

### Motivation

The intestinal epithelium provides crucial barrier, transport and homeostatic functions. These requirements lead it to undergo constant repair and regeneration, and dysfunctions can result in pathologies such as tumorigenesis [1–7]. While much work has been carried out on estimating the rate parameters in the intestine and other epithelia [1, 8–10], attempts to interpret these experimental data using mechanistic modelling remain inconclusive (see e.g. [11–14]). A key issue in drawing reliable conclusions is the lack of consistent and reproducible frameworks for comparing models representing conjectured biological mechanisms, both to each other and to experimental data.

This challenge goes in both directions: using experimental data (taken to be ‘true’) to parameterise and test mathematical or computational formalisations of mechanistic theories, and using these models (taken to be ‘true’) to predict, interpret and question experimental results. Both experimental measurements and mathematical models are subject to uncertainty, and we hence need systematic ways of quantifying these uncertainties and attributing them to the appropriate sources. Furthermore, establishing a common framework for analysing experimental results, formulating mechanistic models and generating new predictions has many potential advantages for enabling interdisciplinary teams to communicate in a common language and efficiently discover and follow promising directions as and when they arise.

### Approach

We address the above issues by developing a best-practice hierarchical Bayesian framework for combining measurements, models and inference procedures, and applying it to a tractable set of experiments targeting mechanisms of repair and regeneration in the intestinal epithelium. These experiments were performed ourselves and are presented in more detail in [15]. The aim of these experiments was to identify how proliferation rates, tissue growth and cellular migration rates are related under healthy, damaged (Ara-C treated) and recovering conditions, and how these relations can be used to identify mechanisms of repair and regeneration.

A notable feature of the Bayesian approach to probabilistic modelling is that all sources of uncertainty are represented via probability distributions, regardless of the source of uncertainty (e.g. physical or epistemic) [16–18]. We will adopt this perspective here, and thus we consider both observations and parameters to be random variables. Within a modelling or theoretical context, uncertainty may be associated with (at least): parameters within a mechanistic model of a biological or physical process, the mechanistic model of the process itself and the measurements of the underlying process. This leads, initially, to postulating a full joint probability distribution for observable, unobservable/unobserved variables, parameters and data.

Another key feature of the Bayesian perspective, of particular interest here, is that it provides a natural way of decomposing such full joint models in a *hierarchical* manner, e.g. by separating out processes occurring on different scales and at different analysis stages. A given set of hierarchical assumptions corresponds to assuming a factorisation of the full joint distribution mentioned above, and gives a more interpretable and tractable starting point.

Our overall factorisation follows that described in [18–21]. This comprises: a ‘measurement model’, which defines the observable (sample) features to be considered reproducible and to what precision they are reproducible (the measurement scale); an underlying ‘process’ model, which captures the key mechanistic hypotheses of spatiotemporal evolution, and a prior parameter model which defines a broad class of *a priori* acceptable possible parameter values.

This hierarchical approach is being increasingly adopted - especially in areas such as environmental and geophysical science [22, 23], ecological modelling [24, 25], as well as in Bayesian statistical modelling and inverse problems more generally [17–21, 26]. In our view, however, many of the advantages of hierarchical Bayesian modelling remain under-appreciated and it offers many opportunities for formulating more unified frameworks for model-data and model-model comparison. Furthermore, we note that a similar hierarchical approach has recently received significant development in the context of the non-Bayesian ‘extended-likelihood’ statistical modelling framework [27–29]. Thus, in our view, many of the benefits of the present approach can be attributed to its hierarchical aspect in particular ([20] also emphasises this point).

As illustration of some of the modelling benefits of the hierarchical approach, we show how both discrete and continuous process models can be derived and related using the hierarchical perspective. We discuss the connection of conditional/hierarchical modelling to the causal modelling literature (see [30–32] for reviews) and illustrate the distinct roles of (Bayesian) predictive distributions vs. parameter distributions for model checking and the assessment of evidence, respectively (see [17, 33–36] for discussion of these distinctions).

### Conclusions

Our hierarchical Bayesian framework incorporates measurement, process and parameter models and facilitates separate checking of these components. This checking indicates, in the present application to the spatiotemporal dynamics of the intestinal epithelium, that much of the observed measurement variability is adequately captured by a simple measurement model. Similarly we find that a relatively simple process model can account for the main spatiotemporal dynamics of interest; however, model checking also identifies a minor misfit in the process model appearing over long time-scales. This motivates possible model improvements: we consider the inclusion of additional finite-cell-size effects in the process model, derived from a discrete process model and a subsequent continuum approximation formulated in terms of conditional probability. This only gives a slightly better qualitative fit to experimental data, however. We instead find that the dominant sources of the long-time misfits are probably due to some other factors - most likely relatively slow, time-varying proliferation rates (e.g. due to circadian rhythms). In summary, a primarily proliferation-driven model appears adequate for predictions over moderate time-scales.

## Materials and methods I: Experimental treatments and data processing

### Homeostasis mouse model

To obtain estimates of intestinal epithelial proliferation and migration rates under normal, homeostatic conditions in healthy mice, we used standard methods of proliferative cell labelling and tracing [1, 8–10, 37–39] (see also [15] for full details). Actively proliferating cells in the intestinal crypts were labelled by single injection of the thymine analogue 5-bromo-2-deoxyuridine (BrdU) and labelled cells detected by immunostaining of intestinal sections collected from different individuals over time. Migration of labelled cells traced from the base of crypts to villus tips was monitored over the course of 96 hours (5760 min). At least 30 strips were analysed per mouse. The figures presented in [15] show that strips were independent and obtained from one-cell thick sections. All strips in which the base of the crypt and the tip of the villus were clearly observed were considered. All sides of the crypt that were visible and connected to entire villi were analysed. There was no arbitrary selection of strips. A typical image from those analysed in [15] is also reproduced in the Supplementary information.

To obtain estimates of intestinal epithelial proliferation and migration rates under normal, homeostatic conditions in healthy mice, we used standard methods of proliferative cell labelling and tracing [1, 8–10, 37–39] (see also [15] for full details). Actively proliferating cells in the intestinal crypts were labelled by single injection of the thymine analogue 5-bromo-2-deoxyuridine (BrdU) and labelled cells detected by immunostaining of intestinal sections collected from different individuals over time. Migration of labelled cells traced from the base of crypts to villus tips was monitored over the course of 96 hours (5760 min). At least 30 strips were analysed per mouse. The figures presented in [15] show that strips were independent and obtained from one-cell thick sections. All strips in which the base of the crypt and the tip of the villus were clearly observed were considered. All sides of the crypt that were visible and connected to entire villi were analysed. There was no arbitrary selection of strips. A typical image from those analysed in [15] is also reproduced in the Supplementary information.

### Blocked-proliferation mouse model

To assess the effects of proliferation inhibition on crypt/villus migration, migrating and proliferating epithelial cells were monitored by double labelling with two thymine analogues (BrdU and IdU), administered sequentially a number of hours apart and subsequently distinguished by specific immunostaining in longitudinal sections of small intestine. Following initial IdU labelling of proliferating cells at t= -17h (-1020 min, relative to Ara-C treatment), mice were then treated with cytosine arabinoside (Ara-C) at 250 mg/kg body weight, a dose reported to inhibit proliferation without causing major crypt cell atrophy (see [15] and references therein for full details). Tissues were collected over 24 hours, with BrdU administered one hour prior to collection to check for residual proliferation. Successful inhibition of proliferation following treatment with Ara-C was confirmed by an absence of BrdU (S-phase) and phospho-Histone H3 (pH3) staining (M-phase) in longitudinal sections of small intestine (again, see [15] for full details).

### Recovering-proliferation mouse model

The above Ara-C treatment effect was observed to last for at least 10h (600 min). Cell proliferation resumed to near normal levels in samples obtained 18h (1080 min) post-Ara-C treatment. We hence considered samples collected at least 10h post-Ara-C treatment as corresponding to ‘recovering-proliferation’ conditions.

### Data processing: reference grid and key observable features

To connect experimental measurements to the models discussed below we specified a reference grid and defined the key features of the data relative to this grid. These key features established an ideal ‘underlying population’ from which samples were considered to be drawn. This also allowed us to construct our hierarchical model in a ‘top-down’ (data-to-parameter) manner, starting from a measurement model.

With reference to Fig 1, we considered the data to consist of a collection of one-dimensional ‘strips’ of cells. These strips ran from the base of the crypt to the tip of the villus, along the so-called ‘crypt-villus’ axis. This corresponds to how strips were collected experimentally, but does not account for possible biases due to ‘angled’ sampling [40, 41]. Each measurement was given a spatial cell location index *i* and a time label *t.* The location index was measured in number of cells along the crypt-villus axis, starting from the crypt base, and hence defined a discrete one-dimensional grid. The two labels *i* and *t* were also combined into a single index parameter *s* ꞉= (*i,t*) when notationally convenient, which then defined a two-dimensional grid of space-time points. A ‘typical’ reference crypt-villus unit was characterised by the two vectors (**L**, **n**), where **L** is the vector of labelled fractions at each grid point and **n** is the vector of number of samples at each grid point. This defines a useful reduction of the system from two spatial dimensions to one.

**Figure 1.**
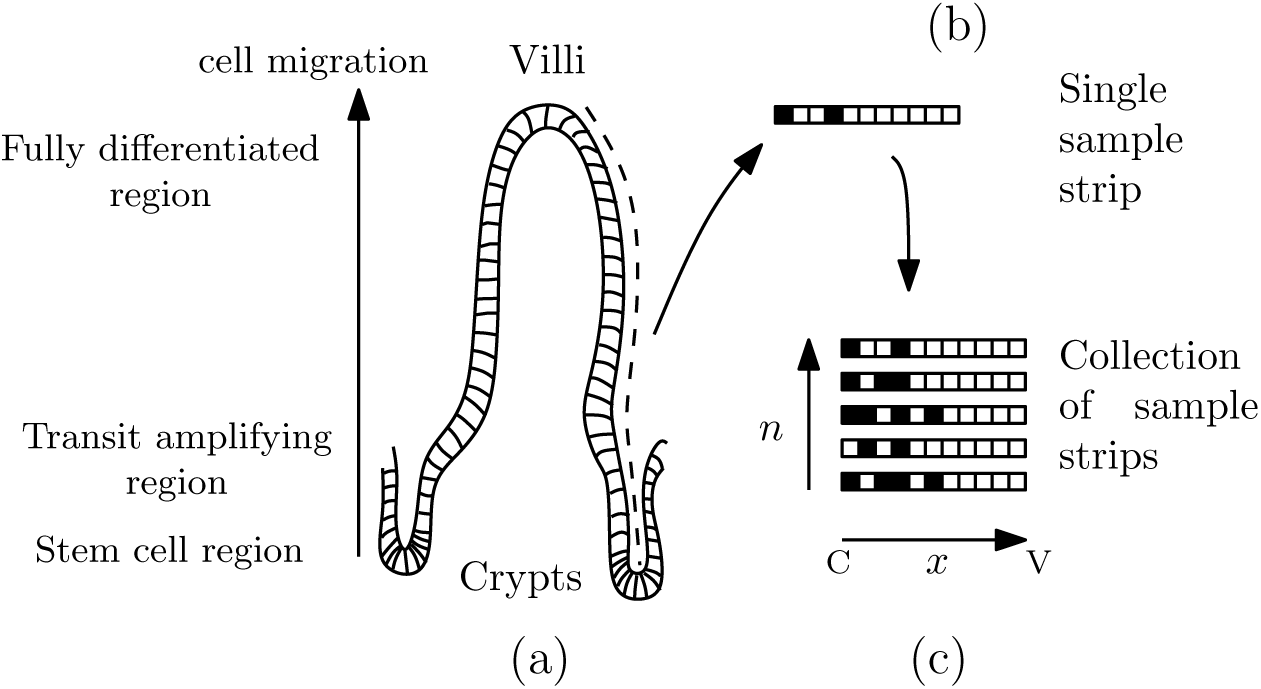
The (a) intestinal epithelium, (b) individual measurements as strips of cells and (c) collection of strips, where ‘C’ and ‘V’ indicated ‘crypt’ and ‘villus’ respectively.

We assumed that each strip was independent of the others as, in general, strips are taken from different crypt-villus units and/or animals after ‘identical preparation’. Thus we did not ever directly possess, for example, measurements of a particular crypt with dimensions given in terms of a certain number of strips. We note, however, that in general the dynamics of strips in a given crypt may be affected by those in the same crypt. We did not consider this additional complexity in the present work, and so this complication should be kept in mind when interpreting the results.

## Materials and methods II: Mathematical framework

Our hierarchical probability model was constructed on the basis of conditional probability assumptions. These allowed us to factor out a measurement model, a mechanistic model and a parameter model.

### Overall hierarchical structure

Our overall model structure consisted of a full joint distribution, conditioned on a given experimental treatment *E* and known sample size vector **n**, decomposed according to

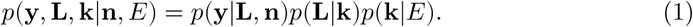

where **k** are the cellular proliferation rates (these are discussed more fully in the Process models and Parameter model sections below). This hierarchical factorisation corresponds to the assumption of conditional independence between the various levels, i.e. *p*(**y**|**L**, **k**, **n**, *E*) = *p*(**y**|**L**, **n**), *p*(**L**|**k**, **n**, *E*) = *p*(**L**|**k**) and *p*(**k**|**n**, *E*) = *p*(**k**|*E*). The first term, *p*(**y**|**L**, **n**) is the *measurement model*; the second term *p*(**L**|**k**) is the underlying *process model*, and the last term *p*(**k**|*E*) is a *prior parameter model.* These are discussed in more detail below.

Notably, a ‘causal’ (structural invariance) assumption [30–32, 42–47] is made by assuming that the experimental treatment condition affects the process parameter **k** but not the structure of the measurement or process models. In general, we suppressed, in our notation, the explicit conditioning on sample size **n**, since it was taken to be fixed and known, as well as the conditioning on *E* (keeping in mind that it only affects **k**).

The assumptions underlying the above factorisation could be checked to some extent. This relied on a distinction between working ‘within’ the model - e.g. parameter estimation assuming the model and factorisation is valid - and working ‘outside’ the model, e.g. checking the validity of the model structural assumptions themselves [17, 35, 36]. This distinction is made in the Results section.

Implicit in the model derivations, discussed below, we used a *deterministic expression of conservation of probability* for the process model, as is typical for such equations [48]. It sufficed for the presentation here to simply replace all functional dependencies on the process variable above with a dependence on the process parameters [18].

### Bayesian framework for predictions and incorporating information from observations

The overall model of the previous section defined our initial ‘generative’ probabilistic model, prior to explicitly incorporating the information from our experimental data. This enabled samples to be drawn from both prior predictive and prior parameter models, in the usual way (see e.g. [17, 49] and the Computational methods section below). In particular, the prior predictive distribution was used in its usual form

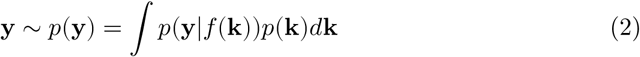

which incorporates the aforementioned deterministic link between a given sample of process parameters and the output process variable, **L** = *f*(**k**). Note that here ~ denotes ‘distributed as’, or more relevantly, ‘samples drawn according to’.

To incorporate new data **y**_0_ we updated the parameters of the model, hence passing to a ‘posterior predictive’ model [17]

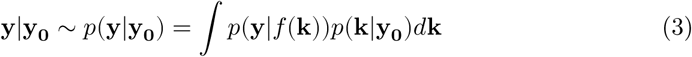

where we used the conditional probability closure assumption *p*(**y**|*f*(**k**), **y_0_**) = *p*(**y**|*f*(**k**)). This closure assumption can be interpreted as maintaining our same mechanistic model despite new observations. This also connects well with current theories of causality as based on ideas of structural invariance [30–32, 42–47].

The logical flow of the updating process we used is depicted in Fig 2. This depicts the ‘forward’ predictive processes as arising from sequences of draws going from ‘lower-level’ to ‘higher-level’ distributions (though this does not correspond directly to the implementation - see Computational methods for specifics). Distributions were updated in the ‘reverse' manner by conditioning at the highest level and propagating the implications of this back down the hierarchy.

**Figure 2.**
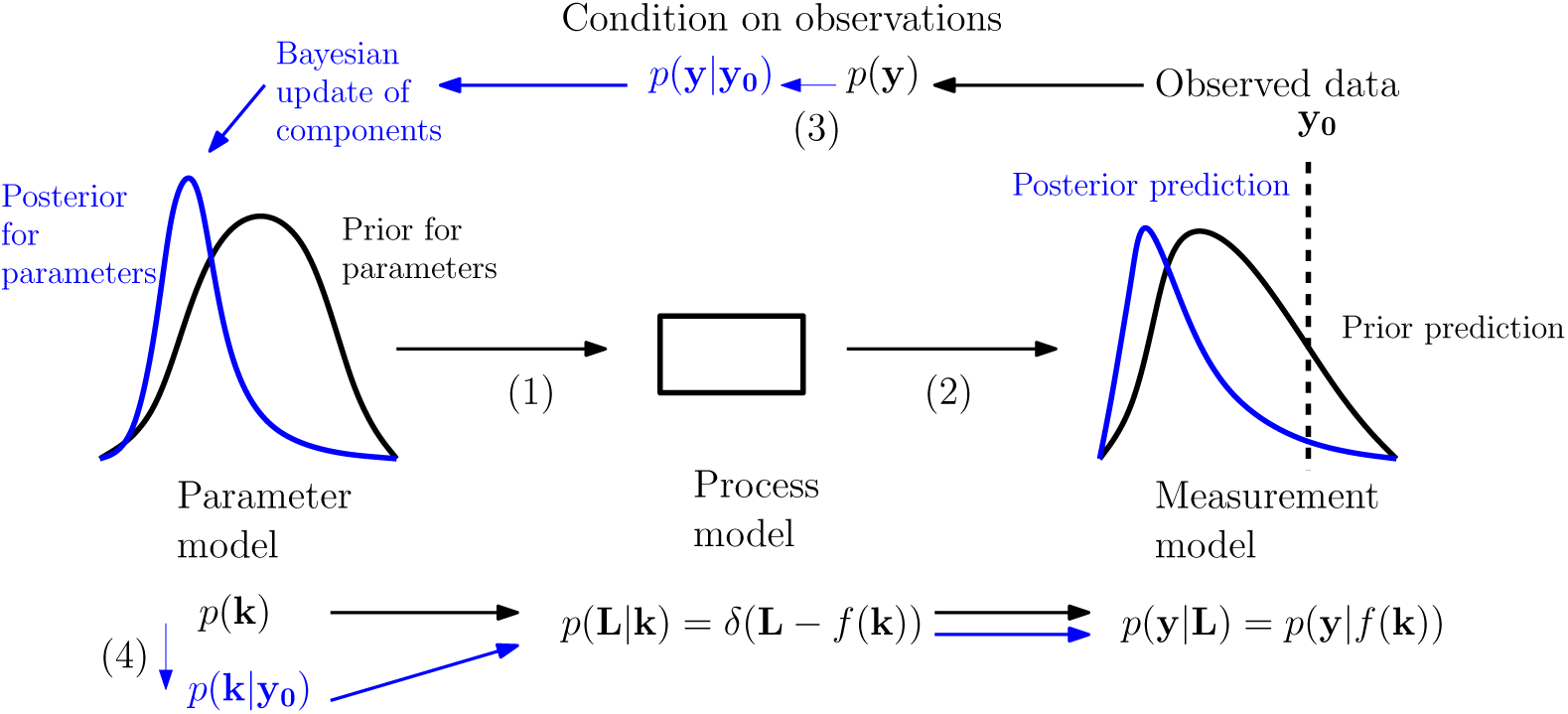
Illustration of the Bayesian predictive and parameter inference processes. Following the arrows (1) to (2) we move from a prior parameter model (left, black) to associated predictive distribution (right, black) via the process and measurement models. Following the arrows (3) to (4) we condition on observed data to obtain a posterior parameter model (left, blue) and associated predictive distribution (right, blue). Our structural assumptions mean that the information gained is represented in updates of the parameter model while the process and measurement models maintain the same form. Modified from [49], which was based on [50].

### Measurement model

The measurement model *p*(**y**|**L**, **n**) component was taken to be a binomial distribution 𝓑 of the form

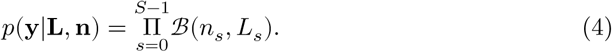

This related our ‘raw’ observable **y**, the vector of counts of labelled cells at each grid point, to ‘ideal characteristics’ of comparison (**L**, **n**),

This was developed as follows. Firstly we assumed that all observations at a given grid point *s* were exchangeable (see [16, 17] for a formal definition and further discussion) conditional on (**L**, **n**). Such exchangeability conditions imply the existence of Bayesian probability models and correspond, in essence, to statistical reduction/symmetry assumptions [16, 17]. We then adopted a slight strengthening [16, 17] of the general exchangeability assumption - which only leads to a pure existence theorem - to an assumption of conditional independence. This assumes that if the true parameters are known at each location then observations can be made independently at those locations.

This latter strengthening assumption is worth noting because it is related to the issue, discussed in the section on experimental methods above, of taking each strip to be independent and the corresponding reduction from two spatial dimensions to one spatial dimension. As such it represents a simplifying approximation and should be kept in mind when interpreting the subsequent results.

We also took the measurement component to be independent of *E* - i.e. treatment was assumed to affect the underlying *process parameters* only (this is discussed in more detail in ‘Overall hierarchical structure’ above, and corresponds to a ‘causal’ assumption).

#### Likelihood and normal approximation

The above defined our measurement component *p*(**y**|**L**, **n**) of the full sampling model for the probability of a set of observed labelled cells **y** in samples of sizes **n** given the vector of modelled underlying labelled fractions **L**. This then defined a likelihood function 𝓛 for this measurement model, which is proportional to the probability given by the sampling model evaluated for the observed data and considered as a function of **L**, i.e.

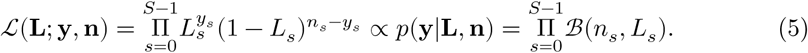

We also used, for interpreting model misfit, the fact that for each *s*, if *n_s_* is sufficiently large and *L_s_* is not too close to 0 or 1 (e.g. *n_s_L_s_* and *n_s_*(1 − *L_s_*) > 5 is typical), then the binomial distribution *B*(*n_s_*, *L_s_*) can be replaced by the normal approximation *N*(*n_s_L_s_*, *n_s_L_s_* ( 1 − *L_s_*)). In this case, denoting the set of all measured labelled fractions through the (useful, but slightly non-standard) notation **y**/**n:***=* (*y*_1_*/n*_1_*,…,y_s_/n_s_*),

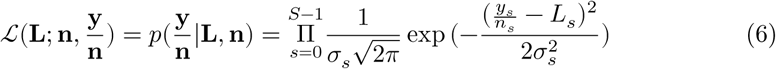

where the standard deviations are given by 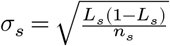. This normal approximation formulation was not used in the model fitting but provided a useful guide for checking model misfit based on residuals.

### Process models

Our process model was developed in a number of stages and considered at different levels of resolution. Firstly, we considered a discrete probabilistic model at the level of our measurement grid defined above. Then we considered two different continuous approximations to this - one excluding explicit cell-scale effects and one including explicit cell-scale effects.

#### Discrete, measurement-grid-level process model

Our basic ‘process’ model described the evolution of the (population) ‘measurement’ probability (labelled fraction) at the scale of the measurement grid. This was derived as follows.

With reference to Fig 1, we considered a collection of one-dimensional ‘strips’ of cells. We used *l_i_* ∈ {0,1} as an indicator variable denoting the occupancy status of site *i* of a given strip. The full state of this strip was given by the vector l = (*l*_0_ *l*_1_, … *l_S–1_*).

We then sought a description of the probabilistic dynamics in terms of a discrete-time Markov chain for the probability distribution of the full state *p*(*l*, *t*) following standard arguments [48, 51].

We began from an explicit joint distribution for the full state and then reduced it to description in terms of the set of ‘single-site’ probability distributions *p*(*l_i_, t*) for each site *i*. This derivation was aided by adopting an explicit notation: the probabilities of occupancy and vacancy at site *i* at time *t* were denoted by *p*(*l_i_*(*t*) = 1) and *p*(*l_i_*(*t*) = 0) respectively. Since *p*(*l_i_*(*t*) = 1) *+p*(*l_i_*(*t*) = 0) = 1 we only needed to consider the probability of occupancy to fully characterise the distribution *p*(*l_i_*(*t*)).

The equation of evolution for this probability was derived by considering conservation of probability in terms of probability fluxes in and out, giving, to first order in Δ*t*

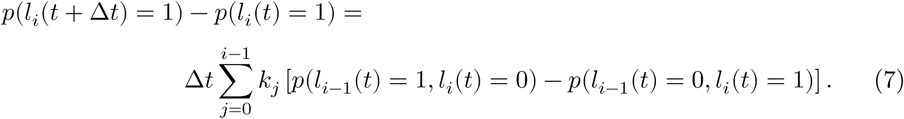

The first term on the right gave a net ‘influx of occupancy probability’ due to a single division event somewhere at site *j < i,* each division event having a probability given by *k_j_* Δ*t.* This flux meant the value of the state variable *l_i_*(*t*) = 0 could be replaced, at the next time step, by the value of *l_i-1_*(*t*) = 1. The second term similarly represented a net ‘outflux of occupancy probability’ due to a division event somewhere at site *j < i.*

Since *l_i_*(*t*) *=* 0 and *l_i_*(*t*) *=* 1 partitioned the event space of *l_i_*(*t*), we could use

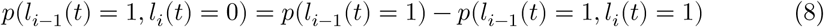

and similarly, since *l_i-1_*(*t*) = 0 and *l_i-1_*(*t*) = 1 partitioned the event space of *l_i-1_*(*t*), we had

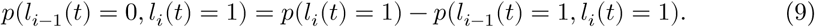

This led to the simplification in terms of only single-site probability distributions

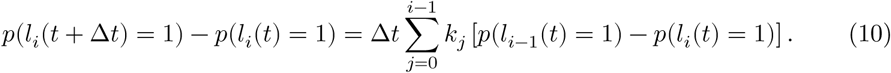

#### Underlying continuous model - zeroth-order approximation

To aid model interpretation and model cross comparisons we introduced a smooth parameter field of underlying labelled fractions *L*(*x*, *t*), defined over a continuous space-time domain. This gave a further idealisation of the ‘underlying population’ from which we envisaged the strips were sampled. This smoothness assumption, while not strictly necessary, meant some model properties could be interpreted in terms of local derivatives; it also reduced arbitrary dependence on discrete grid features, aiding future comparisons with off-lattice and/or continuum models (see [49] for a review of various model types).

To derive the continuous approximation we first defined the position *x* as a continuous coordinate passing through the discrete cell indices. For example *x =* 0 denoted the coordinate of the cell labelled ‘0’ (base of the crypt), while *x =* 0.5 was the location halfway between the cell labelled ‘0’ and that labelled 1’. Sample locations consisting of space-time pairs were denoted by *s =* (*x_s_*, *t_s_*). Then, for sample locations (*i*, *t*) corresponding to cell indices and arbitrary times, we matched the discrete model and continuous model using

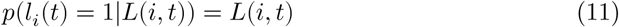

i.e. *L*(*i*, *t*) served as the parameter for a single measurement modelled as a Bernoulli trial at that sample location (as in the above Measurement model section).

Next, the discrete dynamics of *p*(*l_i_*(*t*) = 1) were ‘transferred’ to the continuous *L*(*x*, *t*) dynamics. In particular, since *L*(*x*, *t*) was taken to be a smooth function, we made the correspondence

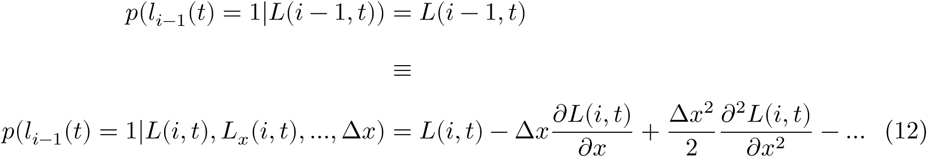

where Δ*x = i* − (*i* − 1) = 1 was the normalised cell length and we also conditioned on knowledge of the spatial derivatives at *i,* 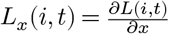 etc. The continuous spatial field effectively interpolated between - i.e. *internal* to - points of the discrete grid, making use of local derivative information. Substituting the above Taylor series, and similar expressions, into the discrete Markov equation 10 led to

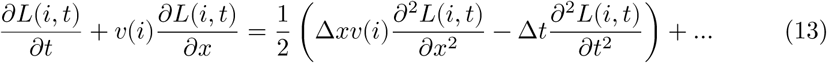

where, for completeness, we also retained higher order terms in Δ*t* for the continuous model. We similarly assumed the existence of smooth functions *k*(*x, t*) and *v*(*x, t*) that satisfied the discrete relations

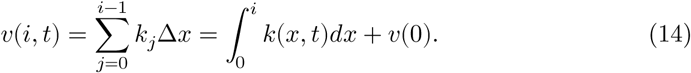

Furthermore, we assumed *k*(*x,t*) = *k*(*x*), *v*(*x,t*) = *v*(*x*) and *v*(0) = 0 in what follows. This assumption is discussed further in the Results section.

We obtained ‘closure’ for the continuous model by keeping only the lowest order terms in both time and space, and further asserting that the equation structure obtained held *for all continuous x* and not just discrete *i* (this could also be motivated by an assumption of grid translation invariance). This gave the advection equation

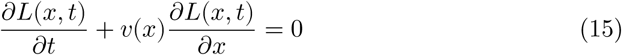

with

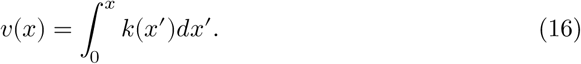

When we incorporated cell death, with discrete rates *d_i_,* this led to the same equations with *k* replaced by *k − d,* where *d*(*x,t*) was defined similarly to *k*(*x,t*). Hence we interpreted *k* in the above as the net cell production rate (which hence could be negative).

The above partial differential equation has an interpretation as the advection of a tracer in an incompressible fluid field with a source, and is sometimes referred to in this context as the ‘colour equation’ [52].

#### Underlying continuous model - higher-order spatial effects

Our ‘zeroth-order’ continuous approximation above was obtained by neglecting all higher-order terms in Δ*x*. We conceived of this as a process of ‘continualisation’ - the reverse process of discretising a continuous equation to obtain a numerical scheme (see e.g. [53] and [52] Section 8.6 for similar ideas). We thus expected that a better continuum approximation could be obtained by retaining higher-order spatial derivatives and hence finite-cell-size effects.

As described below, retaining the higher-order spatial derivative naturally gave rise to a Fokker-Planck equation containing a diffusion term [48]. Equations of this (and similar) form have been derived before, also based on continuous approximations to discrete master equations (e.g. [54–57] also contain similar ideas; [49] gives additional references).

To derive this higher-order approximation we reconsidered the expansion in 13. We again neglected all terms of order Δ*t,* but here retained the next order spatial derivative leading to

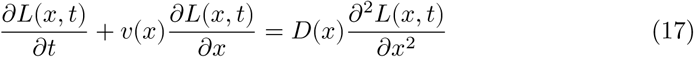

where *D*(*x*) = (*1/2*)Δ*xv*(*x*).

Retaining the second spatial derivative hence amounted to accounting for spatial effects due to finite cell sizes. We first evaluated our original ‘zeroth-order’ (advection) model against our data, but also examined the extent to which higher-order spatial terms such as those considered above could account for any misfits.

### Parameter model

Since we adopted a Bayesian perspective in this work we required a parameter prior model to express additional modelling assumptions ([17] provides an applied account of the role of priors in Bayesian inference, while [36] presents a more philosophical perspective).

Candidate proliferation profiles, varying with cell locations, were represented as realisations from a prior given in terms of a discretised random field (a random vector) **k** of length *m =* 5, modelled as a multivariate Gaussian *N*(*μ,* **C**) with joint distribution

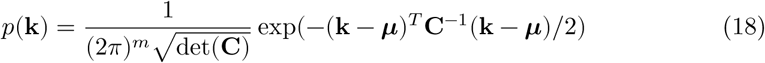

characterised by its mean vector *μ* and covariance matrix **C**. This parameter prior constrained the variability of the spatially varying parameter field *a priori* to help avoid unphysical solutions.

The covariance matrix was first decomposed into a standard deviation matrix given by the outer (tensor) product of the standard deviation vector for each variable, **S** = ***σσ^T^***, and correlation matrix **R**. These multiply element-wise to give *C_ij_* = *S_ij_R_ij_* (no summation). We then adopted the common, equivalent, representation **C** = **DRD** where **D** is a diagonal matrix with diagonal entries *D_ii_* = *σ_i_.*

This decomposition of the covariance matrix into separate parts was adopted because it we felt it presented a clearer picture of how the smoothness and magnitude of variations are controlled via off-diagonal and diagonal terms, respectively, in addition to the mean response. We also varied these prior assumptions to explore the solution dependence on parameter variability (and, as discussed below, our code is made available for further sensitivity tests).

We took the correlation matrix **R** to have the squared-exponential (Gaussian) correlation function 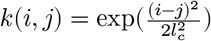, where *l_c_* is a parameter controlling the characteristic length-scale of the correlations in terms of number of indices of **k**. This characteristic length scale gives the number of **k** indices over which the correlation function decays to 1/*e*. This allowed us to control the ‘smoothness’ of the realisations from the **k** prior, in the sense that as *l_c_* is increased the values **k**_*i*_ and **k**_*j*_ tend to be more similar.

The matrix **R** was generated by evaluating this correlation function at discrete locations along the crypt-villus axis. This discretisation was chosen to be coarser than the measurement grid and gave a variation somewhat similar to compartment-style regions of proliferation activity. This corresponded to assuming that the cell-type and associated proliferation rates varied on a coarser scale than individual cells, and was thus somewhat similar to a compartment-style assumption [58, 59], though the resulting proliferation rate function is defined for all values of the finer, individual-cell scale *x.* The parameter *l_c_* could also be interpreted as a ‘parameter correlation length’ for the proliferation rates, a measure of the number of parameters - or number of ‘compartments’ - over which the correlations decay. We considered correlation lengths of 1-2 parameters.

We found it most informative to visualise realisations of the whole function from the resulting prior rather than simply give the individual parameters/matrices separately ([18] discusses this visualisation approach to priors in more detail). These are hence discussed and displayed in more detail in the Results section below.

### Computational methods

#### Implementation of MCMC sampling and Bayesian updating

To implement the updating from prior to posterior parameter distributions, given measurements, we used Monte Carlo Markov Chain (MCMC) sampling (see [60] for a comprehensive reference). In particular, we used the (open source) Python package *emcee* (http://dan.iel.fm/emcee/) which implements an ‘affine-invariant ensemble sampler’ MCMC algorithm and has been applied in particular to astrophysics problems (see [61] for details). Given samples from the resulting prior and posterior parameter distributions, respectively, prior and posterior predictive distributions were obtained by forward simulation of the process model described below. We note that each candidate proliferation rate vector **k** is connected to the measurements **y** via the latent vector **L**; since this step is deterministic, however, no additional sampling steps were required for the process model component.

#### Differential equations

For the results in all sections other than the final results section in which we include higher-order spatial effects, we solved the differential equation model using the *PyCLAW* [62, 63] Python interface to the *CLAWPACK* [64] set of solvers for hyperbolic PDEs. We adapted a Riemann solver for the colour equation available from the Riemann solver repository (https://github.com/clawpack/riemann). For testing the inclusion of higher-order spatial effects (thus changing the class of our equations from hyperbolic to parabolic) we used the Python finite-volume solver *FiPy* [65].

#### Data and source code availability

Our code is available in the form of a Jupyter Notebook

(http://ipython.org/notebook.html) in the Supplementary Information. To run these we used the Anaconda distribution of Python

(https://store.continuum.io/cshop/anaconda) which is a (free) distribution bundling a number of scientific Python tools. Any additional Python packages and instructions which may be required are listed at the beginning of our Jupyter Notebook.

### Interpretation of statistical evidence

We have described above how mechanistic or causal assumptions relate to assumptions of structural invariance under different scenarios. In order to interpret the results that follow, however, we also required an interpretation of the ‘statistical evidence’ that a set of measurements provided about parameter values within a fixed model structure. This proved a surprisingly controversial topic and we encountered continuing debate about fundamental principles and definitions of statistical evidence [35, 66–69].

Following our conditional modelling approach, we decided to adopt the simple - yet quite generally applicable - principle of evidence based on conditional probability: if we observe *b* and *p*(*a|b*) *> p*(*a*) then we have evidence for *a.* A ‘gold-standard’ theory of statistical evidence starting from this premise has been developed and defended recently by Evans in a series of papers (summarised in [35]). Besides simplicity, a nice feature of this approach, that we used below, is that it can be applied both to prior and posterior predictive distribution comparisons such as 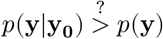, as well as to prior and posterior parameter distribution comparisons such as 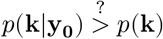 This approach is not without criticism, however (again, see [35, 66–69] for an entry point to the ongoing debates).

Another notable feature of the interpretation of statistical evidence that we adopted below is that we emphasised the visual comparison of various prior and posterior distributions, rather than adopting arbitrary numerical standards ([18] advocates a similar ‘movie strategy’ for the interpretation of statistical evidence and inference procedures, [17, 36, 70, 71] similarly emphasise the benefits of graphical visualisation methods in statistics).

## Results

### Parameter inference under homeostatic (healthy) conditions

Fig 3 illustrates the process of updating from realisations of the prior distributions of the proliferation and velocity fields to realisations of their posterior (post-data) distributions. The left-hand side of the figure shows simulations from the prior distribution for proliferation field (top) and realisations from the induced distribution for the velocity field (bottom), respectively. The right-hand side shows the corresponding simulations after the prior parameter distribution has been updated to a posterior parameter distribution. The prior-to-posterior parameter estimation was carried out using the MCMC sampling approach described above with *t =* 120 min (2 h) as an initial condition and *t =* 360 min (6 h) and 600 min (10 h) as given data. The initial condition for the underlying labelled fraction was determined by fitting a smoothing spline to the data. The prior distribution for the proliferation field shown in Fig 3 incorporated a weak mean trend in net proliferation rates, rising from the crypt base to the mid-crypt before falling exponentially to zero over the last few parameter regions post-crypt end, and a parameter correlation length of 1. These assumptions can be relaxed/varied with little effect, though typically a non-zero parameter correlation length and a shut-off in proliferation after the crypt end produce more stable (well-identified) estimates. As mentioned, the code is available for use and so these assumptions are able to be varied by future researchers. Additional visualisations of the parameter inferences are also provided in the Supplementary information.

**Figure 3.**
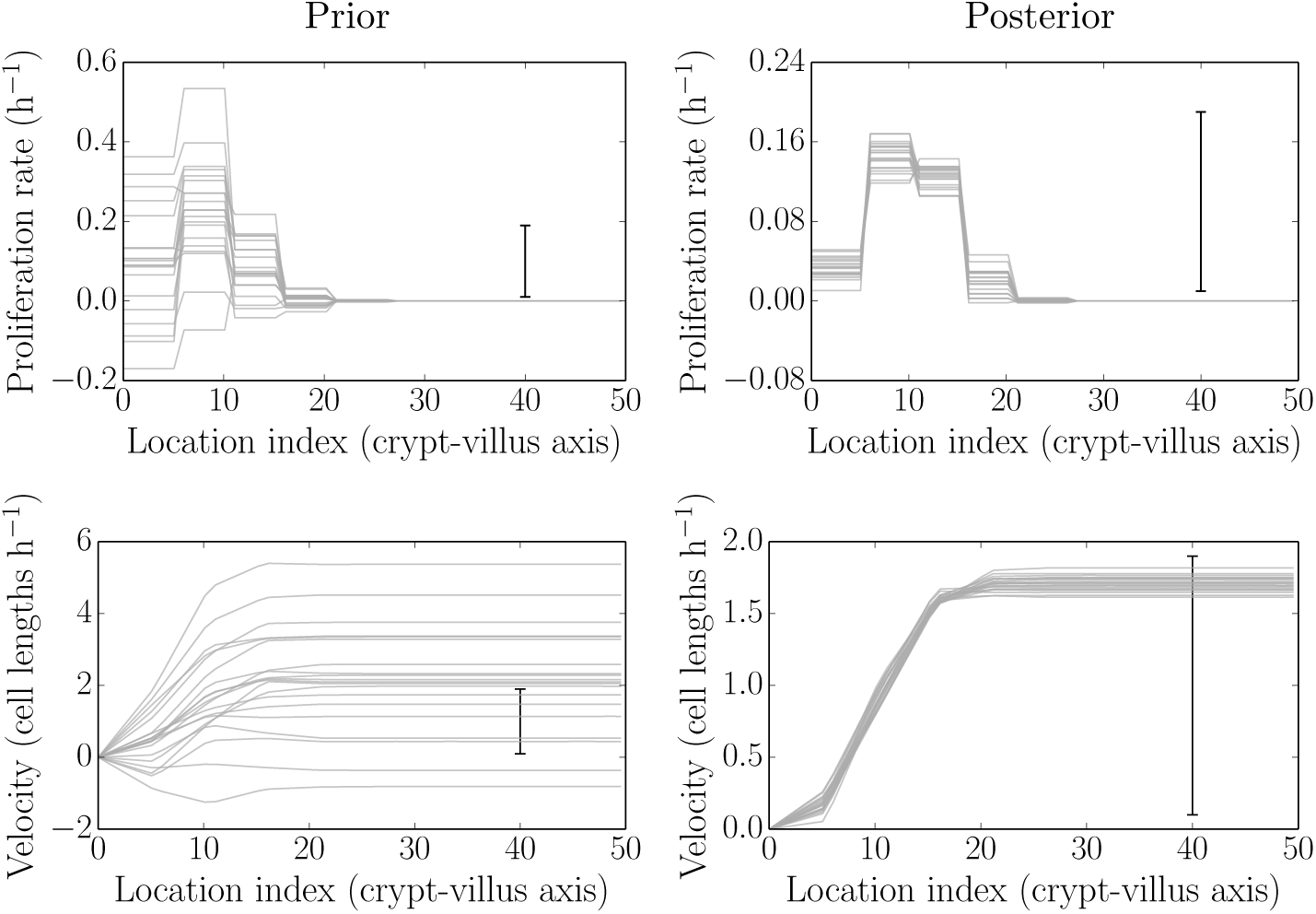
Simulated realisations from the prior (left) and posterior (right) distributions for proliferation profiles (top) and velocities (bottom). After data are obtained the posterior distributions are much more tightly-constrained, and are picking out biologically plausible results (see main text).

### Parameter inference for blocked proliferation conditions

Fig 4 is the same as Fig 3 described in the previous section, but this time under treatment by Ara-C. The previous results from the baseline case are shown in grey, while the new results under Ara-C treatment are shown in blue. Here 1140 min (19 h post IdU labelling, 2 h post Ara-C treatment) was used as the initial condition and 1500 min (25 h post IdU labelling, 8 h post Ara-C treatment) used for fitting. The intermediate time 1260 min (21 h post IdU labelling, 4 h post Ara-C treatment) and later time 1620 min (27 h post IdU labelling, 10 h post Ara-C treatment) were used as out-of-sample comparisons (see later).

**Figure 4.**
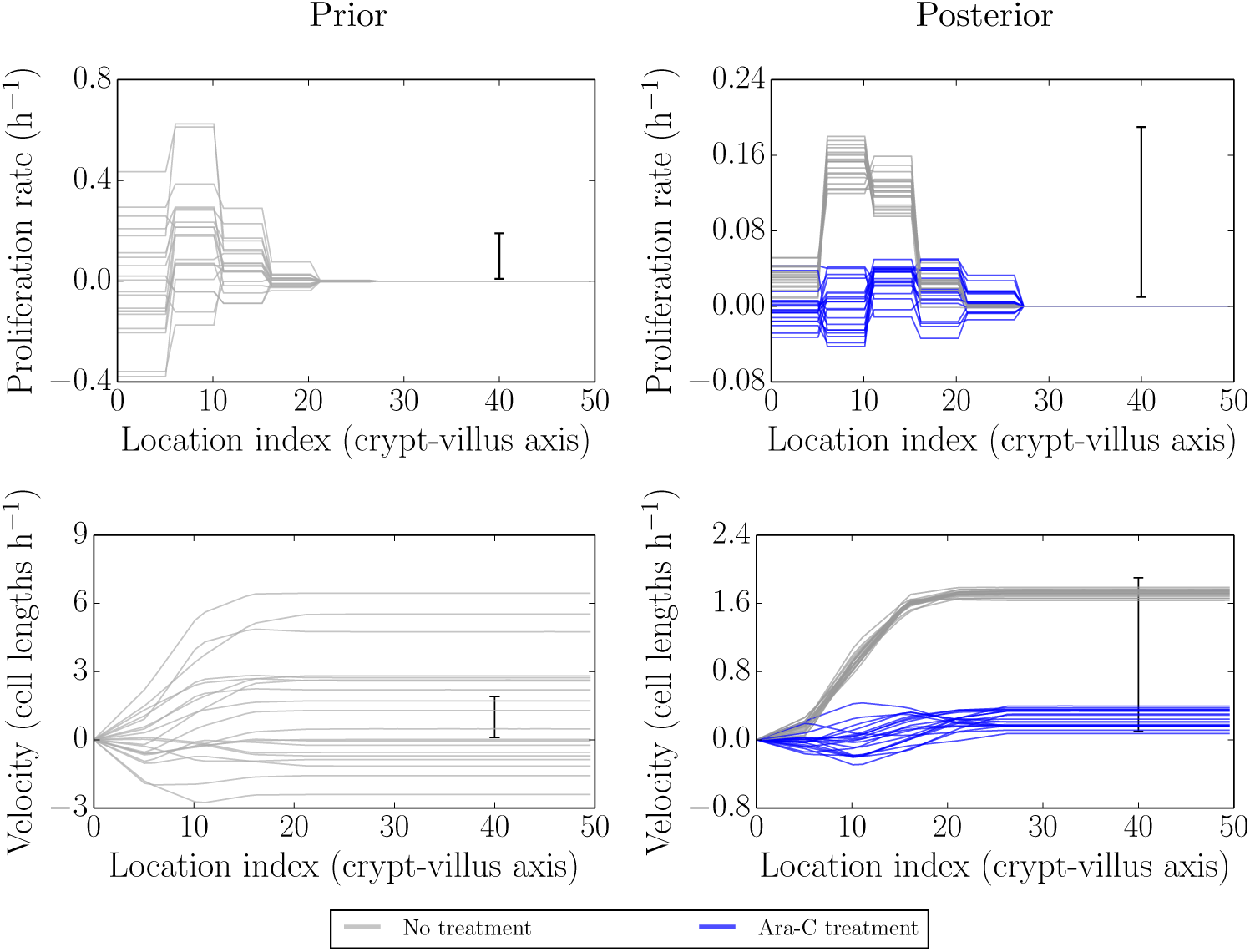
Simulated realisations from the prior (left) and posterior (right) distributions for proliferation profiles (top) and velocities (bottom) under Ara-C treatment (blue) as compared to no treatment (grey). The velocities are reduced to near zero, as are the proliferation rates, though the latter are noisier.

As can be seen, there is a clear inhibition of proliferation and an even clearer effect on the migration (growthf velocity. The underlying parameter results are clearly more variable than those in the baseline case. This may indicate, for example, greater parameter underdetermination and/or inconsistency of the model. This is not surprising as we expect all the proliferation parameters to be reduced to similar (low) values and hence the parameters become less distinguishable.

To add additional stability to the results we can attempt to reduce underdetermination in the parameters by increasing the parameter correlation length and inducing an effectively more ‘lumped’ representation of the parameter field (since values tend to stick together more). Doing this removed the more extreme negative net proliferation in the posterior profile, however it still allowed for small amounts of negative net proliferation/velocity (the available Jupyter notebook can be used to explore various prior assumptions).

Again, the need to introduce more stability is likely due to some combination of the limitations of resolution, a consequence of trying to fit the data too closely, or an indication of model inadequacies. In particular, under inhibited-proliferation conditions the effective number of parameters would be expected to be reduced. When fitting the full model, with largely independent parameters for each region, it is to be expected that some additional regularisation would be required for greater stability.

### Parameter inference for recovering proliferation conditions

Ara-C is metabolised between 10-12 h post-treatment. The two times considered here, 1620 min and 2520 min, correspond to 10 h and 25 h post Ara-C treatment, respectively, i.e to the end of the effect and after the resumption of proliferation. Hence, to check for the recovery of proliferation, we fitted the model using 1620 min as the initial condition and 2520 min as the final time.

Fig 5 is the same as Fig 3 and Fig 4 described in the previous sections, but this time after/during recovering from treatment by Ara-C. The previous results from the baseline case are shown in grey, while the new results following recovery from Ara-C treatment are shown in blue. Here 1620 min (27 h post IdU labelling, 10 h post Ara-C treatment) was used as the initial condition and 2520 min (42 h post IdU labelling, 25 h post Ara-C treatment) used for fitting. We did not make additional out-of-sample comparisons in this case, though in-sample posterior predictive checks were still carried out (see later).

**Figure 5.**
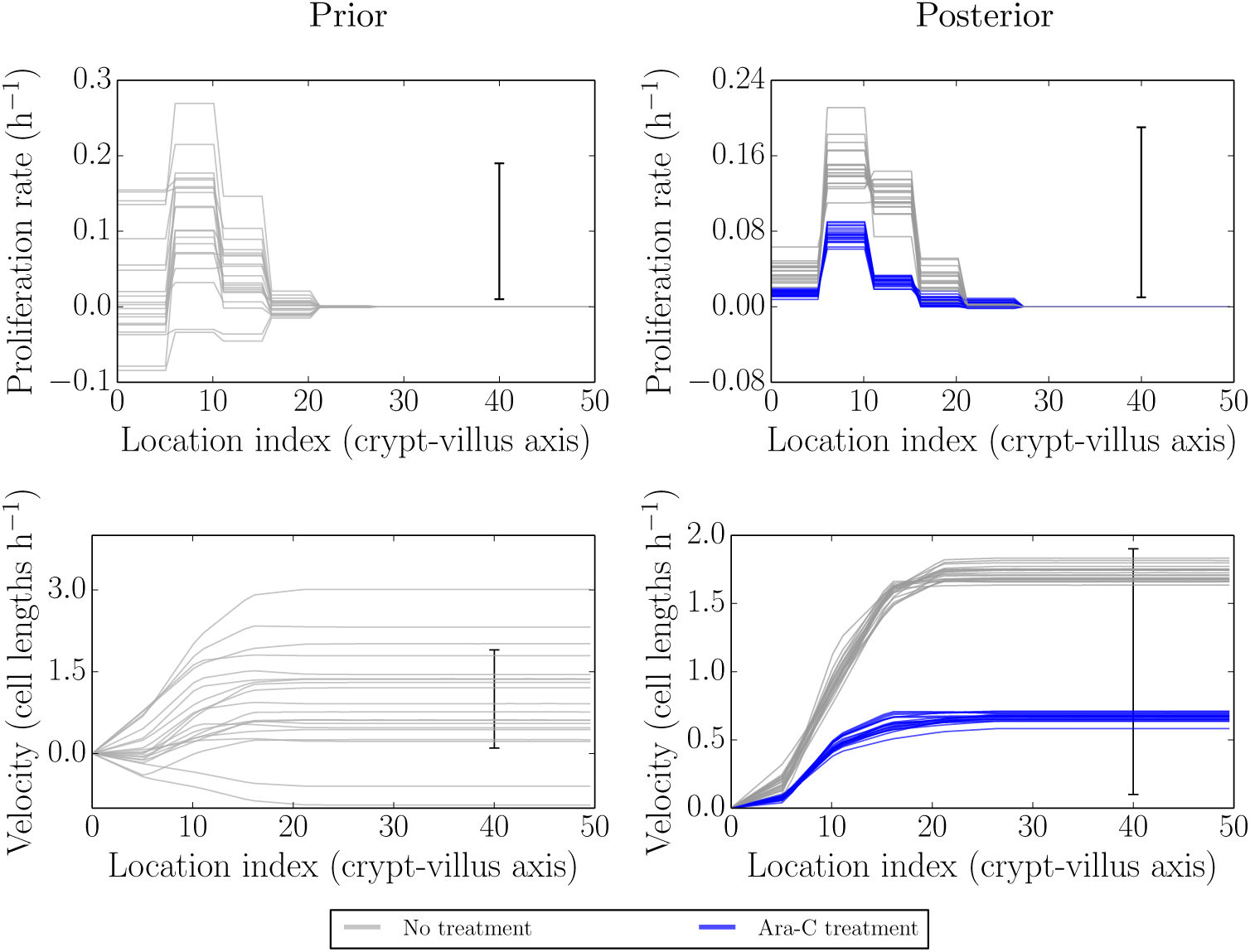
Simulated realisations from the prior (left) and posterior (right) distributions for proliferation profiles (top) and velocities (bottom) after recovery from Ara-C treatment (blue) as compared to no treatment (grey). The velocities and proliferation rates show a clear recovery towards healthy conditions, though not to the full level.

Here the proliferation and velocity profiles indicate that proliferation has resumed, as expected. The rates of proliferation appear to be lower than under fully healthy conditions, however, perhaps due to incomplete recovery (the initial condition being right at the beginning of the recovery period). The timing of the recovery of proliferation and the well-identified proliferation and velocity profiles inferred give no indication that any other mechanism is required to account for these data, however.

### Predictive checks under homeostatic (healthy) conditions

Fig 6 illustrates simulations from the predictive distributions corresponding to the prior and posterior parameter distributions of Fig 3. This enables a first self-consistency check - i.e. can the model re-simulate data similar to the data to which it was fitted [17, 70]. If this is the case then we can (provisionally) trust the parameter estimates in the previous figure; if this was not the case then the parameter estimates would be unreliable, no matter how well-determined they seem. Here the model appears to adequately replicate the data used for fitting.

**Figure 6.**
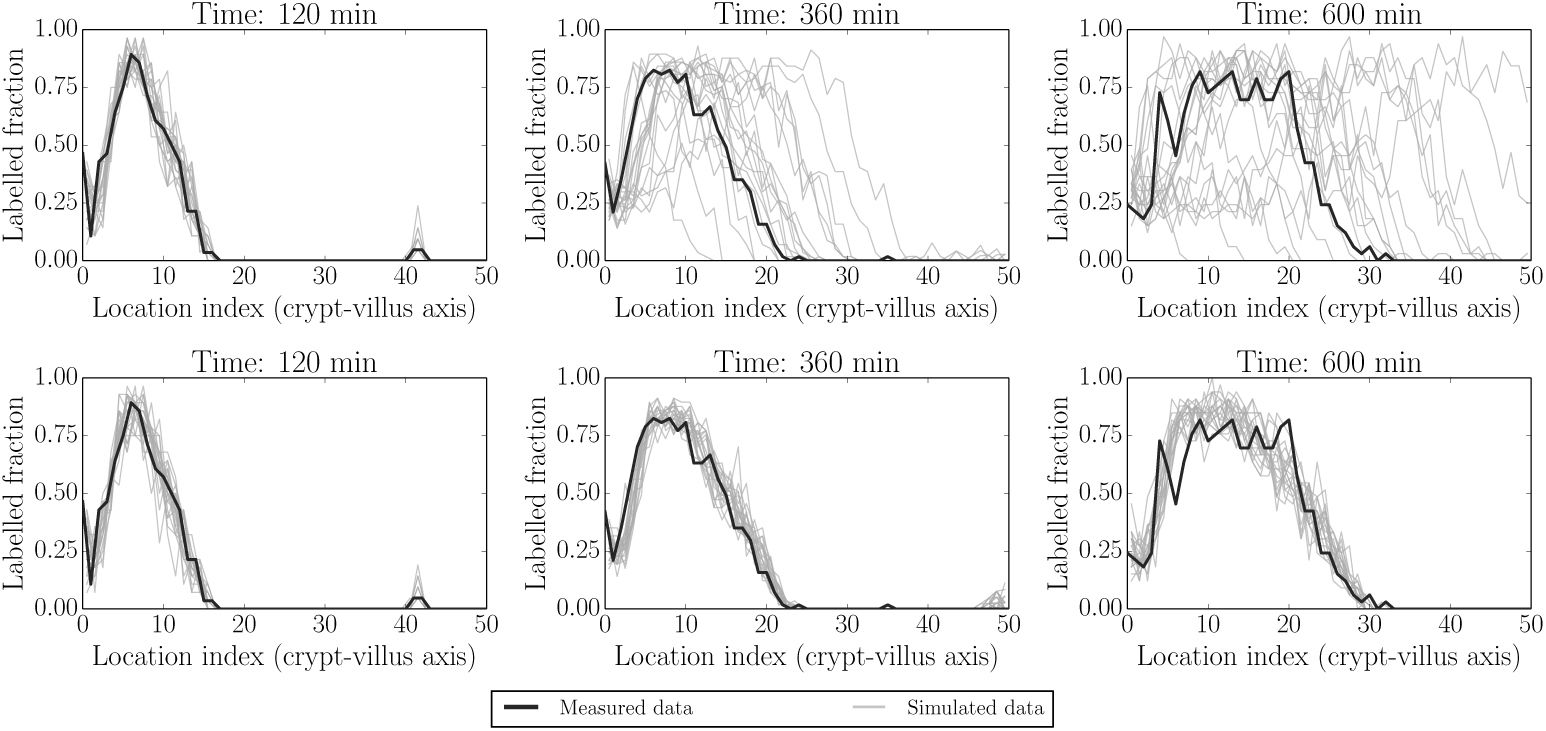
Simulated realisations from prior (top) and posterior (bottom) predictive distributions (grey) for label data at fitted times (120 min, 360 min and 600 min i.e. 2 h, 6 h and 10 h). Actual data are indicated by black lines. Again the posterior distributions are much more constrained than the prior distributions, representing the gain in information from collecting (and fitting to) experimental data. The first profile in each panel is held as a constant initial condition in this example.

Fig 7 and Fig 8 illustrate two additional ways of visualising replicated datasets. The former visualises the label profile along the crypt-villus axis at the future unfitted/out-of-sample time 1080 min (18 h), while the latter visualises both fitted (120 min/2 h, 360 min/6 h and 600 min/10 h) and unfitted/out-of-sample (1080 min/18 h) predictions plotted in the characteristic plane (*t*, *x*) in which the slopes along lines of constant colour should be inversely proportional to the migration velocities at that point, due to the (hyperbolic) nature of our ‘colour’ equation (see e.g. [52]). We have interpolated between the dotted grid lines. These figures, in combination with Fig 6 above, indicate that the model is capable of reliably reproducing the data to which it was fitted, as well as predicting key features of unfitted datasets such as the rate of movement of the front. On the other hand, there is clearly a greater misfit with the predicted rather than fitted data. In order to locate the possible source of misfit we considered various model residuals and error terms - see ‘Locating model misfit’ below.

**Figure 7.**
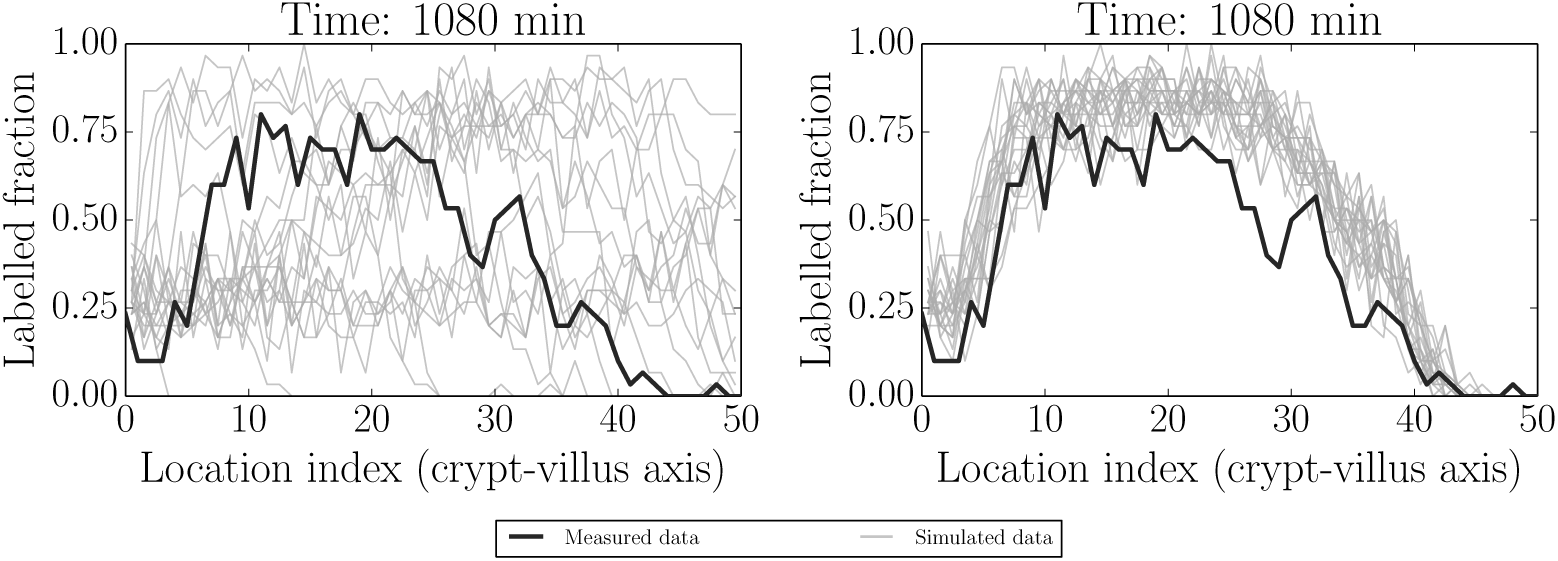
Simulated realisations from prior (left) and posterior (right) predictive distributions (grey) for label data at the unfitted (out-of-sample) time 1080 min (18 h). Actual data are indicated by black lines. The model appears to give reasonable predictions capturing the main effects, but there is also clearly some misfit to be explored further.

**Figure 8.**
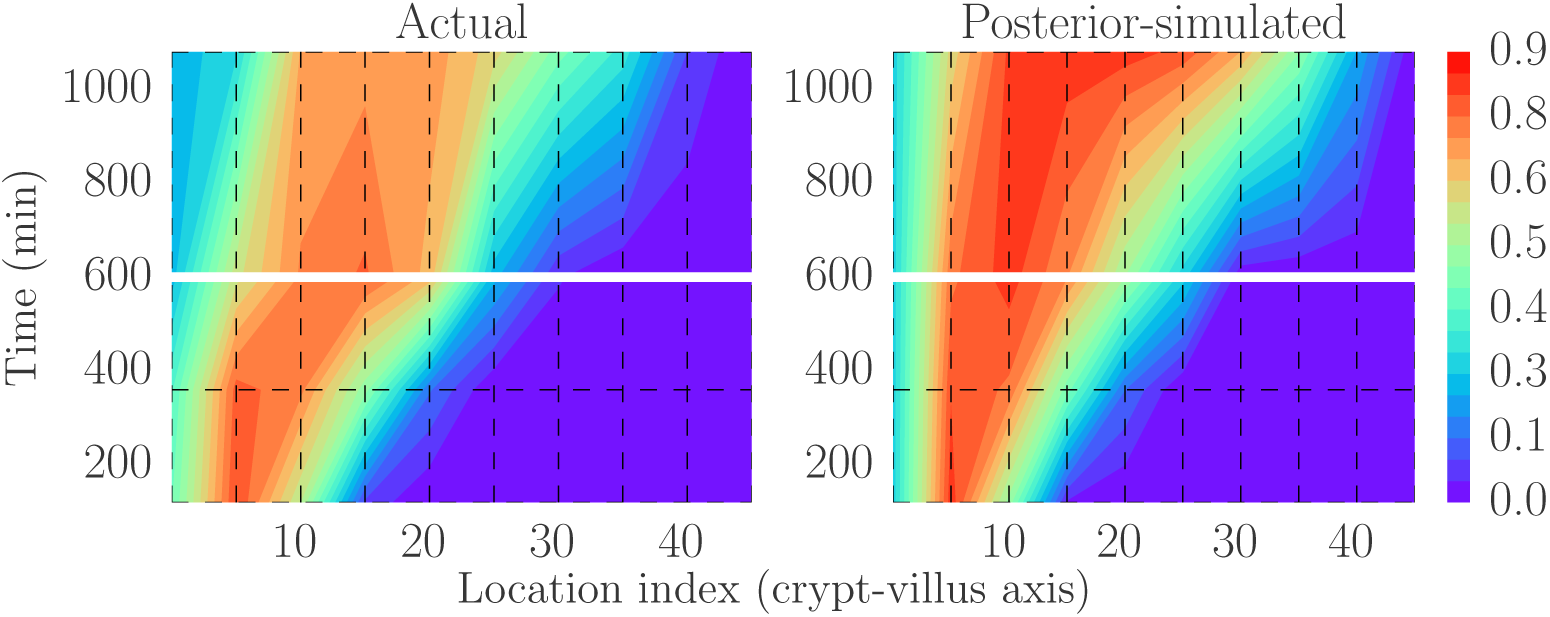
Actual (smoothed) data (left, black box) and one replication based on the model (right; plotting the latent/measurement-error-free process) as visualised in the characteristic plane. This has been discretised and interpolated between the dotted lines to facilitate fair but coarse-grained comparisons. The model structure implies that there should be lines of constant colour tracing out curves with slopes inversely proportional to the migration velocities at that point. The model again captures a number of these key qualitative features, but also fits less well for the out-of-sample (above the horizontal gap at 600 min/10 h) data. There is little variability in the latent model process and so only one replication is shown.

### Predictive checks under blocked proliferation conditions

Here 1140 min (19 h; post IdU labelling) was used as the initial condition and 1500 min (25 h) used for fitting. 1260 min (21 h) and 1620 min (27 h) were used as out-of-sample (non-fitted) comparisons. Fig 9 is analogous to Fig 6 in the healthy case. In general all of the features up to 1620 min (27 h) in Fig 9, and for both fitted and predicted times, appear to be reasonably well captured. The fit at 1620 min is generally good, but perhaps worse than the other cases. This could be due to both errors in longer-time predictions and to the beginning of proliferation recovery. We explore both longer-time misfits and recovering proliferation conditions in what follows.

**Figure 9.**
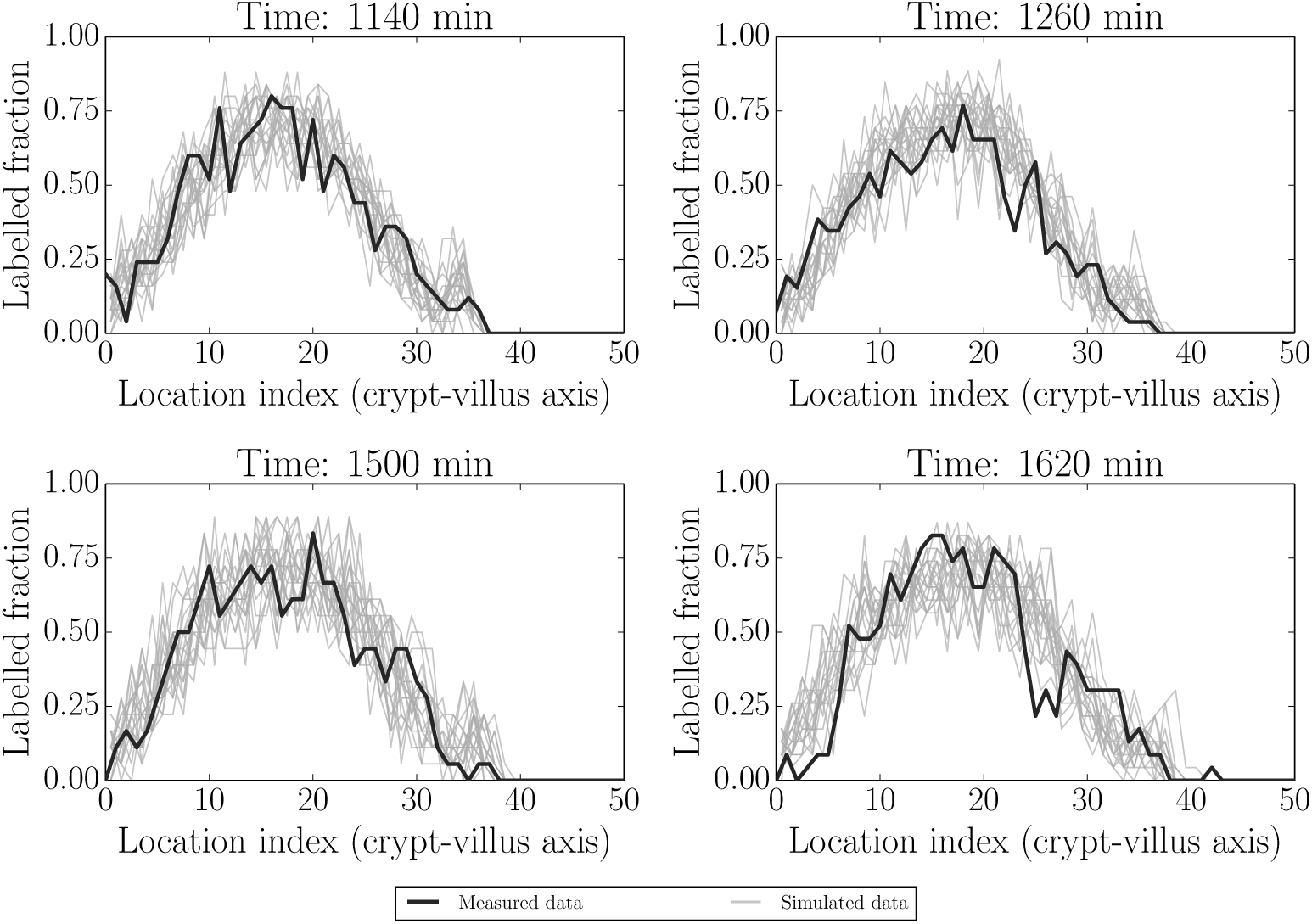
Simulated realisations from posterior predictive distributions (grey) for label data at 1140 min (initial condition), 1500 min (fitted time) and at two out-of-sample/unfitted times (1260 and 1620 min). The posterior distributions appear to adequately capture the actual label data (black).

**Figure 10.**
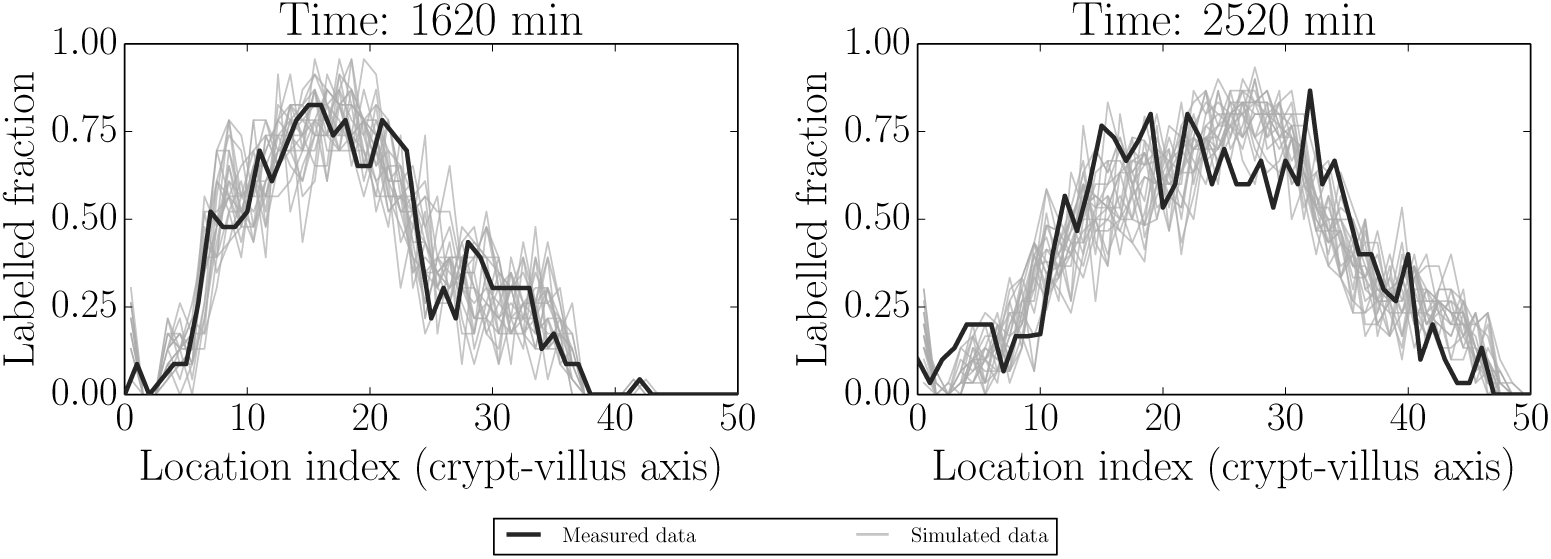
Simulated realisations from posterior predictive distributions (grey) for label data at 1620 min (initial condition) and 2520 min (fitted). These indicate that proliferation has resumed, consistent with the time taken to metabolise Ara-C - see the main text for more detail.

### Predictive checks under recovering proliferation conditions

As discussed above, Ara-C is metabolised between 10-12 h post-treatment. The two times considered here, 1620 min and 2520 min, correspond to 10 h and 25 h post

Ara-C treatment, respectively, i.e to the end of the effect and after the resumption of proliferation.

Again, as expected, the label has resumed movement in concert with the resumption in proliferation. The model appears to fit reasonably well.

### Locating model misfit

While the zeroth-order model behaves essentially as desired under experimental perturbation, and is likely capturing the essential features of interest, we observed some some minor model misfit. We used posterior predictive checks to unpick the contributions of the various model parts and determine the source(s) of misfit. This in turn motivated potential model improvements. These checks were carried out under baseline (healthy) conditions as we were more confident of the experimental effects under this scenario, but they can equally be carried out for the other datasets. Note, however, that time-varying effects are not expected to be as relevant under conditions of inhibited proliferation.

Fig 11 shows the following checks: measurement error as determined by subtracting a smoothed spline from the observed data (dark line) and comparing these to the results obtained by subtracting the process model from the simulated data (panels 1-4, moving left-to-right and top-to-bottom, showing fitted - 120 min/2 h, 360 min/6 h and 600 min/10 h - and unfitted/out-of-sample - 1080 min/18 h - times). This presentation follows the noise-checking approach in [72], as well as the general recommendations given in [17, 70]. Reliable interpretation of these as ‘true’ measurement residuals depends on the validity of the normal approximation 6 since these expressions are not directly interpretable in terms of the discrete binomial model (see e.g. [17, 70]). These are also visualised in terms of the corresponding cumulative distributions in the middle panel (panel 5, following as above). Panels 6-9 show the differences between the underlying process model and the smoothed spline fitted to the data. As can be seen, the measurement model appears approximately valid at all times, while the process model appears to have non-zero error for the 1080 min sample. We consider this in more detail next.

**Figure 11.**
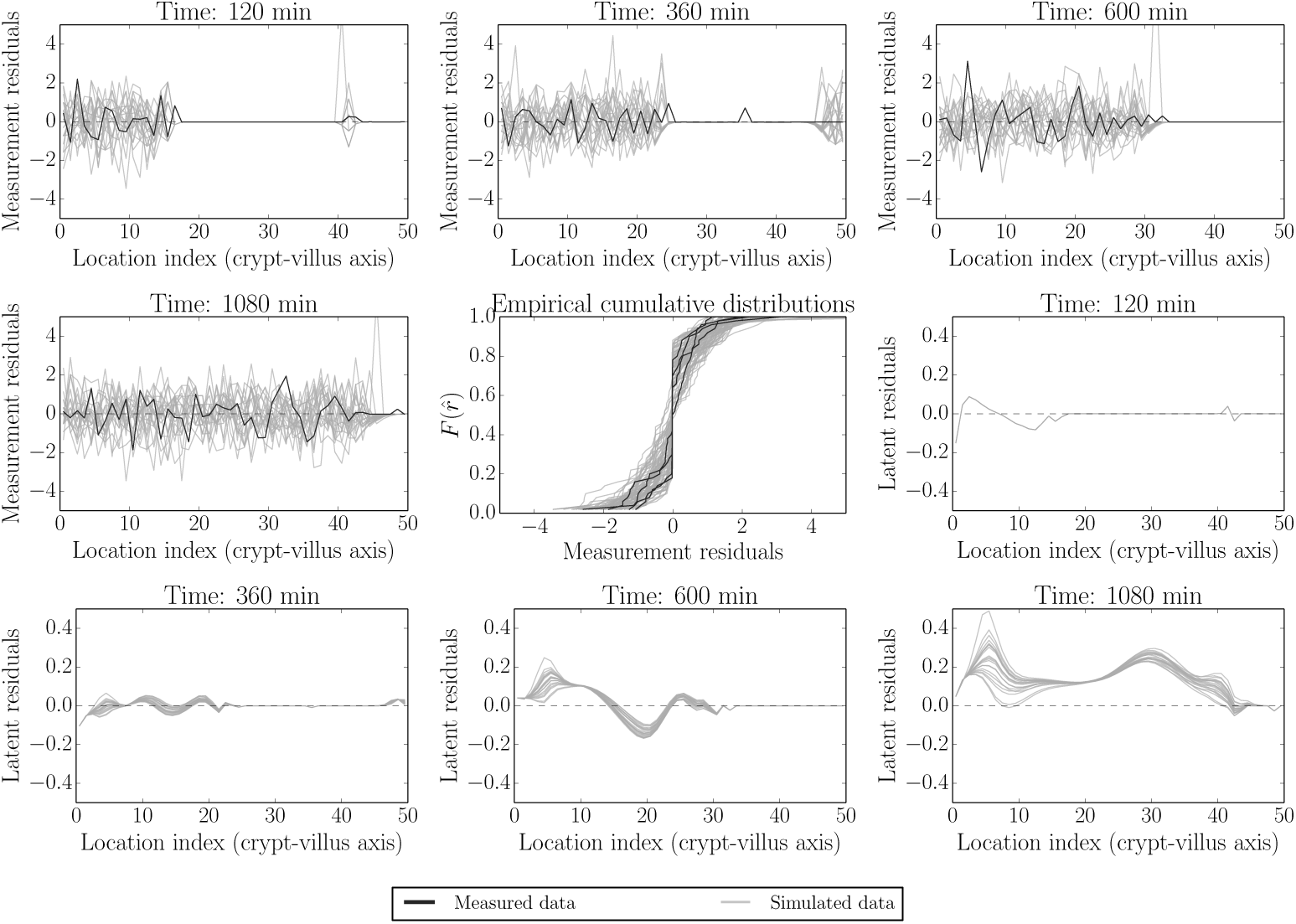
Model and data residual components. Panels 1-4, moving left-to-right and top-to-bottom, shows measurement error as determined by subtracting a smoothed spline from the observed data (dark line) and comparing this to the results obtained by subtracting the process model for fitted - 120, 360 and 600 mins - and unfitted/out-of-sample - 1080 min - times from the realised data (grey). These measurement error distributions are also visualised in terms of the corresponding cumulative distributions in the middle panel (panel 5, following as above. Black - actual data, grey - model simulations). Panels 6-9 show the differences between realisations of the underlying process model and the smoothed spline fitted to the data. As can be seen across panels, the measurement model appears approximately valid at all times, while the process model appears to have non-zero error for the 1080 min sample. This observation is discussed in the text.

### Possible model improvement and robustness - higher-order spatial effects

As discussed in the process model section above, the presence of cellular structure in the epithelial tissue means that higher-order spatial effects could be present. One way of deciding whether these are important is to consider the extent to which these may account for the minor misfit identified above, as opposed to other factors such as time-varying proliferation rates. To do this we considered both uniform percentage reductions of the original parameter estimates (approximating time-varying rates) and the inclusion of higher-order spatial terms.

Fig 12 gives an idea of the qualitative differences induced by including the higher-order spatial terms and those that could be induced by time-varying proliferation rates. This figure is based on the (healthy) 1080 min (18 h) data in which we found some indication of a process model error.

**Figure 12.**
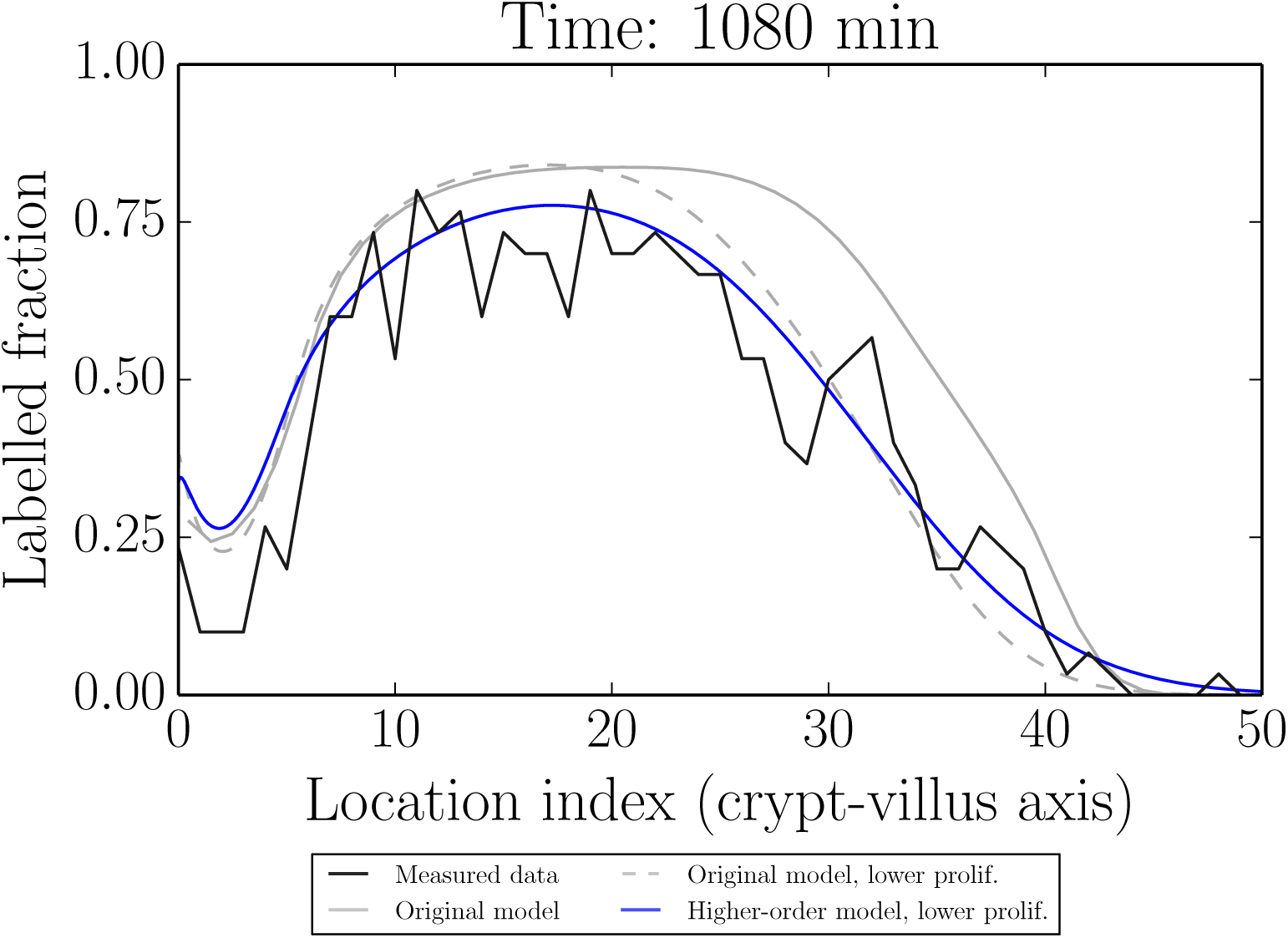
Comparison of the modified process model which includes higher-order spatial terms (blue) to the original model (grey, dashed), both at lowered proliferation rates (decreased 20%), which is required for a better fit to the data. The original model at the original fitted proliferation rates is also shown (grey, solid). Although the model with higher-order spatial terms gives a better qualitative fit to the data for the same proliferation rates, it is clear that the dominant cause of misfit is better attributed to (time) varying proliferation rates (in the context of the present set of models).

We see that while the higher-order model appears to give a slightly better qualitative fit to the data, both the higher-order and lower-order models require similar reductions of the parameter values to quantitatively improve the fit to our out-of-sample data. The reduced parameter values shown in Fig 12 correspond to a reduction of 20%, which was chosen visually as a reduction accounting for the bulk of the misfit.

Thus the key (yet relatively small) difference between the model and out-of-sample data is likely due to an effect other than finite-cell sizes; in this case it is likely due to time-variation in parameter values due to circadian rhythms (we have assumed steady-state parameter values). Other potential factors include label dilution or an unmodelled mixing phenomenon in the full two-dimensional case. We note however that these effects are small and appear to be important primarily for predicting much further ahead in time than the fitted data and the steady-state parameter assumption is likely valid for reasonable time intervals. This means that the more easily interpretable original model may be sufficient for many purposes.

## Discussion

Understanding the complicated dynamics of the intestinal epithelium requires an interdisciplinary approach involving experimental measurements, mathematical and computational modelling, and statistical quantification of uncertainties. While a diverse range of mathematical models have been constructed for epithelial cell and tissue dynamics (reviewed in [58, 59, 73–75]), from compartment models to individual-based models to continuum models, we lack consistent and reproducible frameworks for comparing models representing conjectured biological mechanisms both to each other and to experimental data (for an overview, see our review [49]). These shortcomings may explain why questions such as the connection between proliferation and migration and its variation under experimental perturbations remain open, despite much investigation [8–14].

The aim of the present work was to acknowledge and confront these difficulties explicitly, and to present some initial constructive steps in establishing such a framework. To do this we carried out new experiments (described more fully in a companion paper [15]) aimed at determining how proliferation rates, tissue growth and cellular migration rates are related in the intestinal epithelium under healthy, damaged (Ara-C treated) and recovering conditions. We performed BrdU/IdU cell-labelling experiments under these respective conditions. In considering how to best process these data and interpret them using mathematical models, we then developed a probabilistic, hierarchical (conditional) framework.

Our hierarchical framework provides a best-practice approach for systematically modelling and understanding the uncertainties that could lead to unreliable mechanistic conclusions - uncertainties in experimental measurement and treatment, difficult-to-compare mathematical models of underlying mechanisms, and unknown or unobserved parameters. Our approach was influenced by recognising the benefits that the hierarchical Bayesian approach has demonstrated in applications across a number of different disciplines (e.g. in environmental and geophysical science as in [22, 23]; ecological modelling as in [24, 25]; and in Bayesian statistical modelling and inverse problems more generally as in [17–21, 26]). We also note that a hierarchical approach can have significant benefits outside the Bayesian framework (see for example the ‘extended likelihood’ approach described in [27–29]).

The hierarchical approach has advantages not only in terms of providing a framework for combining disparate sources of uncertainty, but also as a framework for facilitating modelling derivations and relating discrete and continuous models. Though the resulting measurement, process and parameter models can or have all been derived by other means, as far as we are aware this particular perspective has not been systematically utilised in the same manner as considered here - at the very least it appears uncommon within the mathematical/systems/computational biology communities. Furthermore, in the main text we noted the connections of our conditional, probabilistic approach for relating discrete and continuous models to similar procedures in the numerical analysis literature. This raises exciting connections to the developing field of probabilistic numerical methods and computing [76].

We also note the connection between the choice of a measurement model as required here (and/or process model error, and following e.g. [18–22, 77]), and the development of approximate sampling and parameter fitting procedures, which are particularly useful for analytically difficult models. A key concern of the latter is the appropriate choice of summary statistics for constructing a ‘synthetic likelihood’ [78] or similarly-modified posterior target for Approximate Bayesian Computation (ABC) [79–81]. This choice determines (implicitly or explicitly) in which ways a given model or set of models can be considered an ‘adequate’ representation of the data, which features are considered to be reproducible and what the associated ‘noise’ structure should be ([71] presents an alternative approach to characterising data features and model adequacy). These issues are crucial in deciding how to model the complexity of epithelial cell and tissue dynamics.

An important next step, as described above, would be to bring more process model types into this framework and to evaluate and compare them under carefully modelled experimental conditions. Extensions incorporating other mechanical and/or cellular-level information (e.g. [11, 12]) into process models would provide a natural next step. Importantly, due to the separation between measurement and process model components, these more complex process models could be incorporated into our present framework simply by replacing our process model component with a new model, while retaining the same measurement model. Of course additional parameters would require additional prior assumptions, and if additional data features were of interest then these would need to be incorporated into a modified measurement model. The benefit of a hierarchical framework is that it offers an explicit guide as to where such modifications should be incorporated.

We also agree with the view expressed by [17] that the cycle, adopted here, of model construction, parameter estimation, (graphical) model checking and model expansion is typically more informative than ‘model selection’ as traditionally understood - especially when this latter activity is based on Bayes factors or other assignments of single numerical quantities to complex models. We generally advocate understanding and comparing models in terms of predictive checks and identifying which features particular models capture well and which they miss. That is, we do not believe that there is much to gain from choosing one model as ‘best’ - rather that we should understand in which respects our models are ‘useful’ [33, 34]. Part of our goal here was to encourage more researchers to think in these terms and point out that the hierarchical approach has the potential to facilitate such analyses for a range of different model types.

As a final methodological point, by making our code and data available, as well as leveraging already-available open-source scientific Python software, we open up our work to other researchers to build on.

In summary, the main results established using the above framework were

- An adequate description of intestinal epithelial dynamics is achievable using a model based on purely proliferation-driven growth
- This model is consistent with healthy, proliferation-inhibited (Ara-C treated) and recovering conditions
- The measurement and process model errors can be reasonably distinguished and checked separately
- This checking indicates that much of the natural variability is directly attributable to the collection process and this process can be modelled in a simple manner
- Possible model errors can also be identified and proposed explanations incorporated and tested within our framework, and thus the proper interpretation of experimental procedures is aided by using an explicit mathematical model and its predictive simulations
- Including finite-cell-size effects gives a slightly better qualitative fit to experimental data, but the dominant sources of the long-time misfits are likely due to some other factor such as (relatively slowly) time-varying proliferation rates (e.g. due to circadian rhythms) or label dilution.

## Acknowledgements

This work was funded by the BBSRC-UK, project numbers BB/K018256/1, BB/K017578/1, BB/K017144/1 and BB/J004529/1 and the EPSRC-UK, project number EP/I017909/1.

## References

1. Wright NA, Alison M (1984) The biology of epithelial cell populations. Oxford University Press, USA

2. Radtke F, Clevers H (2005) Self-renewal and cancer of the gut: Two sides of a coin. Science 307:1904–1909

3. Reuss L (2010) Epithelial transport. Comprehensive physiology

4. van der Flier LG, Clevers H (2009) Stem cells, self-renewal, and differentiation in the intestinal epithelium. Annu Rev Physiol 71:241–260

5. Turner JR (2009) Intestinal mucosal barrier function in health and disease. Nat Rev Immunol 9:799–809

6. Marchiando AM, Graham WV, Turner JR (2010) Epithelial barriers in homeostasis and disease. Annu Rev Pathol 5:119–144

7. Barker N (2014) Adult intestinal stem cells: Critical drivers of epithelial homeostasis and regeneration. Nat Rev Mol Cell Biol 15:19–33

8. Kaur P, Potten CS (1986) Cell migration velocities in the crypts of the small intestine after cytotoxic insult are not dependent on mitotic activity. Cell Tissue Kinet 19:601–610

9. Kaur P, Potten CS (1986) Circadian variation in migration velocity in small intestinal epithelium. Cell Tissue Kinet 19:591–599

10. Tsubouchi S (1983) Theoretical implications for cell migration through the crypt and the villus of labelling studies conducted at each position within the crypt. Cell Tissue Kinet 16:441–456

11. Dunn S-J, Näthke IS, Osborne JM (2013) Computational models reveal a passive mechanism for cell migration in the crypt. PLoS One 8:e80516

12. Meineke FA, Potten CS, Loeffler M (2001) Cell migration and organization in the intestinal crypt using a lattice-free model. Cell Prolif 34:253–266

13. Loeffler M, Stein R, Wichmann HE, Potten CS, Kaur P, Chwalinski S (1986) Intestinal cell proliferation. I. A comprehensive model of steady-state proliferation in the crypt. Cell Tissue Kinet 19:627–645

14. Loeffler M, Potten CS, Paulus U, Glatzer J, Chwalinski S (1988) Intestinal crypt proliferation. II. Computer modelling of mitotic index data provides further evidence for lateral and vertical cell migration in the absence of mitotic activity. Cell Tissue Kinet 21:247–258

15. Parker A, Maclaren OJ, Fletcher AG, Muraro D, Kreuzaler PA, Byrne HM, Maini PK, Watson AJM, Pin C (2016) Cell proliferation within small intestinal crypts is the principal driving force for cell migration on villi. The FASEB Journal; published ahead of print October 20, 2016, doi:10.1096/fj.201601002

16. Bernardo JM, Smith AFM (2009) Bayesian theory. John Wiley & Sons

17. Gelman A, Carlin JB, Stern HS, Dunson DB, Vehtari A, Rubin DB (2013) Bayesian data analysis, third edition. Taylor & Francis

18. Tarantola A (2005) Inverse problem theory and methods for model parameter estimation. SIAM

19. Berliner LM (1996) Hierarchical Bayesian time series models. In: Maximum entropy and Bayesian methods. Springer Netherlands, pp 15–22

20. Cressie N, Wikle CK (2011) Statistics for spatio-temporal data. John Wiley & Sons

21. Wikle CK (2015) Modern perspectives on statistics for spatio-temporal data. WIREs Comput Stat 7:86–98

22. Berliner LM (2003) Physical-statistical modeling in geophysics. J. Geophys. Res. 108:

23. Wikle CK (2003) Hierarchical models in environmental science. Int Stat Rev 71:181–199

24. Cressie N, Calder CA, Clark JS, Ver Hoef JM, Wikle CK (2009) Accounting for uncertainty in ecological analysis: The strengths and limitations of hierarchical statistical modeling. Ecol Appl 19:553–570

25. Ogle K (2009) Hierarchical Bayesian statistics: Merging experimental and modeling approaches in ecology. Ecol Appl 19:577–581

26. Blei DM (2014) Build, compute, critique, repeat: Data analysis with latent variable models. Annual Review of Statistics and Its Application 1:203–232

27. Pawitan Y (2001) In all likelihood: Statistical modelling and inference using likelihood. OUP Oxford

28. Pawitan Y, Lee Y (2016) Wallet game: Probability, likelihood and extended likelihood. Am Stat 0:1–7

29. Lee Y, Nelder JA, Pawitan Y (2006) Generalized linear models with random effects: Unified analysis via h-likelihood. CRC Press

30. Dawid AP (2002) Influence diagrams for causal modelling and inference. Int Stat Rev 70:161–189

31. Dawid PA (2010) Seeing and doing: The Pearlian synthesis. In: Rina Dechter, Hector Geffner, and Joseph Y. Halpern(ed) Heuristics, probability and causality: A tribute to judea pearl. College Publications London, pp 309–325

32. Dawid PA (2010) Beware of the DAG! NIPS Causality: Objectives and Assessment 6:59–86

33. Box GEP (1976) Science and statistics. J Am Stat Assoc 71:791–799

34. Box GEP (1980) Sampling and Bayes’ inference in scientific modelling and robustness. J R Stat Soc Ser A 143:383–430

35. Evans M (2015) Measuring statistical evidence using relative belief. CRC Press

36. Gelman A, Shalizi CR (2013) Philosophy and the practice of Bayesian statistics. Br J Math Stat Psychol 66:8–38

37. Kozar S, Morrissey E, Nicholson AM, Heijden M van der, Zecchini HI, Kemp R, Tavaré S, Vermeulen L, Winton DJ (2013) Continuous clonal labeling reveals small numbers of functional stem cells in intestinal crypts and adenomas. Cell Stem Cell 13:626–633

38. Vermeulen L, Morrissey E, Heijden M van der, Nicholson AM, Sottoriva A, Buczacki S, Kemp R, Tavaré S, Winton DJ (2013) Defining stem cell dynamics in models of intestinal tumor initiation. Science 342:995–998

39. Lopez-Garcia C, Klein AM, Simons BD, Winton DJ (2010) Intestinal stem cell replacement follows a pattern of neutral drift. Science 330:822–825

40. Meineke FA, Potten CS, Loeffler M (2001) Cell migration and organization in the intestinal crypt using a lattice-free model. Cell Prolif 34:253–266

41. Potten CS, Roberts SA, Chwalinski S, Loeffler M, Paulus U (1988) Scoring mitotic activity in longitudinal sections of crypts of the small intestine. Cell Tissue Kinet 21:231–246

42. Pearl J (2009) Causality. Cambridge University Press

43. Pearl J (2009) Causal inference in statistics: An overview. Stat Surv 3:96–146

44. Dawid PA (1979) Conditional independence in statistical theory. J R Stat Soc Series B Stat Methodol 41:1–31

45. Woodward J (2003) Making things happen: A theory of causal explanation. Oxford University Press

46. Woodward J (1997) Explanation, invariance, and intervention. Philos Sci 64:S26–S41

47. Spirtes P, Glymour CN, Scheines R (2000) Causation, prediction, and search. MIT Press

48. Van Kampen NG (1992) Stochastic processes in physics and chemistry. Elsevier

49. Maclaren OJ, Byrne HM, Fletcher AG, Maini PK (2015) Models, measurement and inference in epithelial tissue dynamics. Math Biosci Eng 12:1321–1340

50. Iglesias M, Stuart A (2014) Inverse problems and uncerrainty quantification. SIAM news (july/August)

51. Wilkinson DJ (2011) Stochastic modelling for systems biology. CRC Press

52. LeVeque RJ (2002) Finite volume methods for hyperbolic problems. Cambridge University Press

53. Askes H, Metrikine AV (2005) Higher-order continua derived from discrete media: Continualisation aspects and boundary conditions. Int J Solids Struct 42:187–202

54. Baker RE, Yates CA, Erban R (2010) From microscopic to macroscopic descriptions of cell migration on growing domains. Bull Math Biol 72:719–762

55. Hywood JD, Hackett-Jones EJ, Landman KA (2013) Modeling biological tissue growth: Discrete to continuum representations. Phys Rev E Stat Nonlin Soft Matter Phys 88:032704

56. Fozard JA, Byrne HM, Jensen OE, King JR (2010) Continuum approximations of individual-based models for epithelial monolayers. Math Med Biol 27:39–74

57. Murray PJ, Edwards CM, Tindall MJ, Maini PK (2009) From a discrete to a continuum model of cell dynamics in one dimension. Phys Rev E Stat Nonlin Soft Matter Phys 80:031912

58. Johnston MD, Edwards CM, Bodmer WF, Maini PK, Chapman SJ (2007) Examples of mathematical modeling: Tales from the crypt. Cell Cycle 6:2106–2112

59. Carulli AJ, Samuelson LC, Schnell S (2014) Unraveling intestinal stem cell behavior with models of crypt dynamics. Integr Biol 6:243–257

60. Robert C, Casella G (2013) Monte carlo statistical methods. Springer Science & Business Media

61. Foreman-Mackey D, Hogg DW, Lang D, Goodman J (2013) Emcee: The MCMC hammer. Publications of the Astronomical Society of the Pacific

62. Ketcheson DI, Mandli K, Ahmadia AJ, Alghamdi A, Luna MQ de, Parsani M, Knepley MG, Emmett M (2012) PyClaw: Accessible, extensible, scalable tools for wave propagation problems. SIAM J Sci Comput 34:C210–C231

63. Mandli KT, Ketcheson DI, others (2014) PyClaw software.

64. Clawpack Development Team (2014) Clawpack software.

65. Guyer JE, Wheeler D, Warren JA (2009) FiPy: Partial differential equations with python. Comput. Sci. Eng.

66. Royall R (1997) Statistical evidence: A likelihood paradigm. CRC Press

67. Mayo DG (2014) On the birnbaum argument for the strong likelihood principle. Stat Sci 29:227–239

68. Evans M (2014) Discussion of “on the birnbaum argument for the strong likelihood principle”. Stat Sci 29:242–246

69. Taper ML, Ponciano JM (2016) Evidential statistics as a statistical modern synthesis to support 21st century science. Popul Ecol 58:9–29

70. Gelman A (2004) Exploratory data analysis for complex models. J Comput Graph Stat 13:755–779

71. Davies PL (2014) Data analysis and approximate models: Model choice, Location-Scale, analysis of variance, nonparametric regression and image analysis. CRC Press

72. Aguilar O, Allmaras M, Bangerth W, Tenorio L (2015) Statistics of parameter estimates: A concrete example. SIAM Rev 57:131–149

73. Kershaw SK, Byrne HM, Gavaghan DJ, Osborne JM (2013) Colorectal cancer through simulation and experiment. IET Syst Biol 7:57–73

74. De Matteis G, Graudenzi A, Antoniotti M (2013) A review of spatial computational models for multi-cellular systems, with regard to intestinal crypts and colorectal cancer development. J Math Biol 66:1409–1462

75. Fletcher AG, Murray PJ, Maini PK (2015) Multiscale modelling of intestinal crypt organization and carcinogenesis. Math Models Methods Appl Sci 25:2563–2585

76. Hennig P, Osborne MA, Girolami M (2015) Probabilistic numerics and uncertainty in computations. Proc Math Phys Eng Sci 471:20150142

77. Mosegaard K, Tarantola A (2002) Probabilistic approach to inverse problems. International Geophysics Series 81:237–268

78. Wood SN (2010) Statistical inference for noisy nonlinear ecological dynamic systems. Nature 466:1102–1104

79. Marin J-M, Pudlo P, Robert CP, Ryder RJ (2012) Approximate Bayesian computational methods. Stat Comput 22:1167–1180

80. Wilkinson RD (2013) Approximate Bayesian computation (ABC) gives exact results under the assumption of model error. Stat Appl Genet Mol Biol 12:129–141

81. Ratmann O, Andrieu C, Wiuf C, Richardson S (2009) Model criticism based on likelihood-free inference, with an application to protein network evolution. Proc Natl Acad Sci U S A 106:10576–10581

